# A Model of Temporal Scaling Correctly Predicts that Weber’s Law is Speed-dependent

**DOI:** 10.1101/159590

**Authors:** Nicholas F. Hardy, Vishwa Goudar, Juan L. Romero-Sosa, Dean V. Buonomano

## Abstract

Timing is fundamental to complex motor behaviors: from tying a knot to playing the piano. A general feature of motor timing is temporal scaling: the ability to produce motor patterns at different speeds. Here we report that temporal scaling is not automatic. After learning to produce a Morse code pattern at one speed, subjects did not accurately generalize to novel speeds. Temporal scaling was also not a general property of a recurrent neural network (RNN) model, however after training across different speeds the model produced robust temporal scaling. The model captured a signature of motor timing—Weber’s law—but predicted that temporal precision increases at faster speeds. A human psychophysics study confirmed this prediction: the standard deviation of responses in absolute time were lower at faster speeds. These results establish that RNNs can account for temporal scaling, and suggest a novel psychophysical principle: the Weber-speed effect.

---

It is increasingly clear that the brain uses different mechanisms and circuits to tell time across different tasks. For example, distinct brain areas are implicated in sensory^1,2^ and motor^3–6^ timing tasks on the scale of hundreds of milliseconds to a few seconds. This “multiple clock” strategy likely evolved because different tasks have distinct computational requirements. For example, judging the duration of a red traffic light relies on the judgement of absolute temporal durations, but tying your shoe and playing the piano rely on the relative timing and order of activation of similar sets of muscles. A general property of these complex forms of motor control is temporal scaling: well-trained motor behaviors can be executed at different speeds. Despite the importance of temporal scaling in the motor domain, basic psychophysical and computational questions remain unaddressed. For example, is temporal scaling intrinsic to motor timing? In other words, once a complex pattern is learned can it be accurately sped-up or down, like choosing the speed at which a movie is played?

The neural mechanisms underlying temporal scaling are unknown in part because the neural mechanisms underlying motor timing itself continue to be debated. Converging evidence from theoretical ^7–9^ and experimental studies suggests that motor timing is encoded in patterns of neural activity, i.e. population clocks^4,5,10–15^. Numerous computational models have been proposed to account for timing^16,17^ but the problem of temporal scaling remains largely unaddressed.

Here we show that under the appropriate training conditions an RNN model exhibits temporal scaling. The model also accounts for a signature feature of motor timing referred to as the scalar property or Weber’s law: the standard deviation of timed responses increases linearly with time^18^. However, the model predicts that the relationship between variance and time is not constant, but dependent on speed. A psychophysical study in which humans produce a complex pattern of taps confirmed this prediction: humans were more precise at producing a response at the same absolute time during a pattern executed at a higher speed.

## RESULTS

It is well established that humans can execute well-trained complex movements such as speaking or playing a musical instrument at different speeds. However, it is not clear if once a complex temporal pattern is learned, whether it can automatically be executed at different speeds. A few studies have examined temporal scaling in humans^19,20^, however, to the best of our knowledge no studies have trained subjects to learn a complex temporal pattern at a single speed, across days, and then examined subject’s ability to reproduce that pattern at faster and slower speeds. We thus first addressed the question of whether temporal scaling is an intrinsic property of motor timing by training subjects on a temporal pattern reproduction task (Online Methods). Over a four-day training period subjects learned to tap out a Morse Code pattern (the word “time”) at a speed of 10 words-per-minute (the duration of a “dot” was 120 ms). The target pattern was composed of six taps and lasted 2.76 s (Fig. 1A).

**Figure 1.**
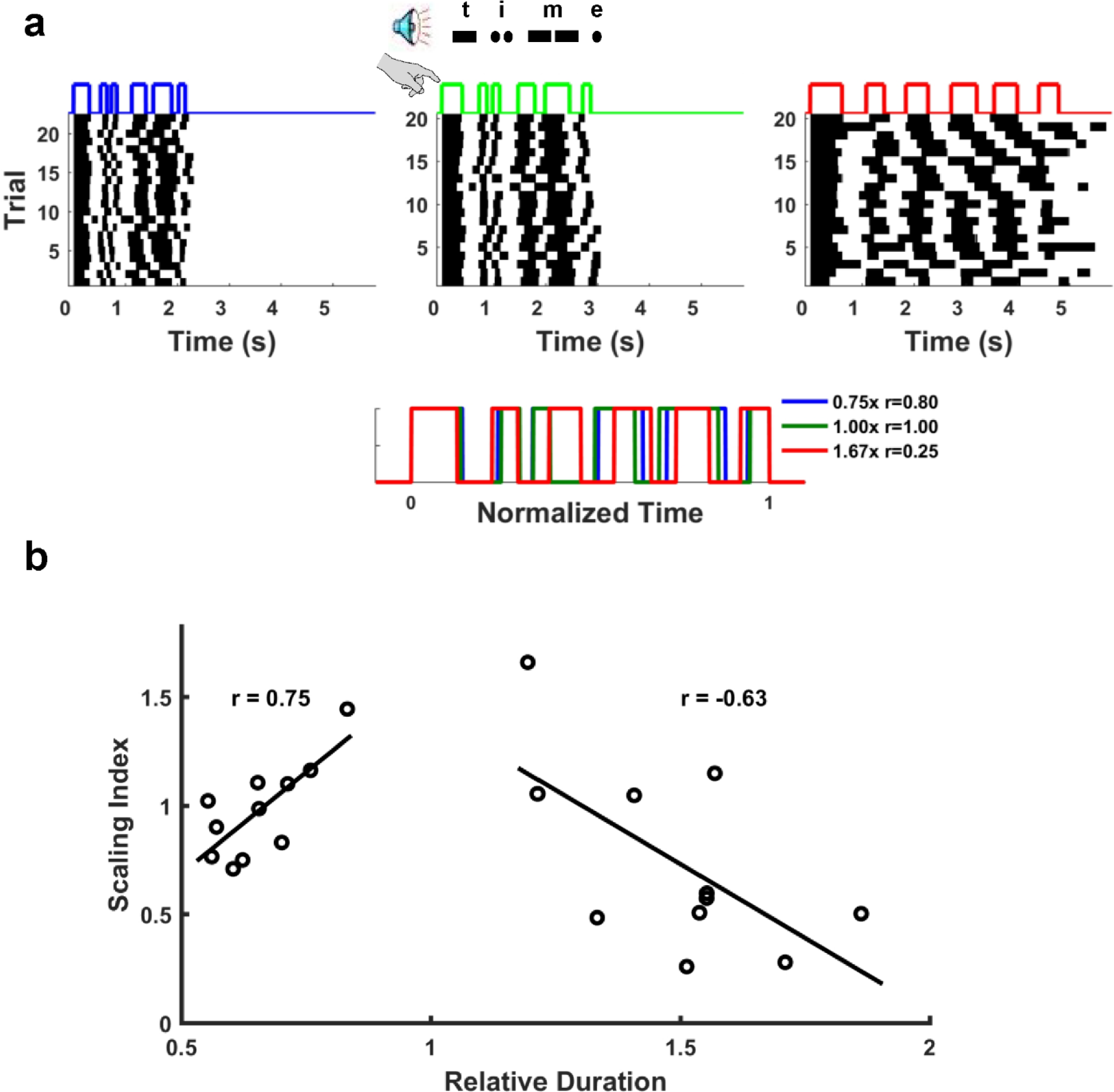
Limited temporal scaling of a learned Morse code pattern. Subjects were trained to tap the Morse code for “time” at ten words per minute over four consecutive days (Online Methods). **a)** On the fifth day, subjects were asked to produce the pattern at three different speeds: twice as fast (2x), normal speed (1x), and twice as slow (0.5x). Bottom: Average of the responses above plotted in normalized time. The legend indicates the produced speed relative to the trained (1x) condition and the correlation of the mean response to the response at trained speed. **b)** The relationship between produced speed and temporal scaling accuracy for all eleven subjects. There was a significant correlation between speed and accuracy for both the fast (r=0.75, p=0.008) and slow (r=-0.63, p=0.038) patterns.

The day after the end of training, subjects were tested on their ability to produce the pattern at the original speed, twice as fast (50% duration), and at half speed (200% duration) under freeform conditions—i.e., they were not cued with the target pattern during this test phase. At the 1x speed subjects were able to reproduce the word “time” in Morse code with a performance score (correlation between the produced and target patterns) of 0.66 ± 0.04. As expected in a freeform condition, few subjects reached the speeds of 2x and 0.5x. Thus we were able to measure how well subjects were able to temporally scale the trained pattern, and whether the accuracy of temporal scaling was related to the magnitude of scaling (speed). We quantified temporal scaling using a scaling index based on the time normalized correlations (Online Methods) between the 1x and 2x and 0.5x patterns (Fig. 1B). For both the fast and slow patterns there was a significant correlation between the scaling index and overall pattern duration (r=0.75, p=0.008; and r=-0.63, p=0.038, respectively). These results confirm that with moderate levels of training, humans are able to speed up or slow down a learned pattern to some extent, but that performance degraded significantly at slower and faster speeds.

### RNN Model of Motor Timing

How can neural circuits generate similar temporal patterns at different speeds? To elucidate the potential mechanisms of temporal scaling, we turned to a RNN model that has previously been shown to robustly generate both simple and complex temporal patterns^8^. The model is composed of randomly connected firing rate units whose initial weights are relativity strong, placing the network in a high-gain regime. In this regime RNNs exhibit complex (high-dimensional) patterns of activity. In theory, this activity can encode time while retaining long-term memory traces on scales much longer than the time constants of the units, but in practice it cannot because the dynamics of the RNN are chaotic^21^. Specifically, chaotic behavior impairs the networks’ computational capacity because network activity patterns are not reproducible across trials. It is possible, however, to tune the recurrent weights to “tame” the chaos while maintaining their complexity (Online Methods). The result is the formation of locally stable trajectories, i.e. dynamic attractors, that robustly encode temporal motor patterns. We first asked whether these RNNs can account for temporal scaling.

One of the most intuitive neural mechanisms of temporal scaling is that the level of external drive to a network changes the speed of its dynamics. Thus, to test whether this class of RNN could account for temporal scaling, we examined the effects of altered input drive. The RNNs in this study receive two independent inputs: one serving as a cue to start a trial and the second as a tonic speed input (*y_SI_*), to modulate the speed of the dynamics. The recurrent units generate motor patterns through synapses onto a single output unit (Fig. 2A).

**Fig 2.**
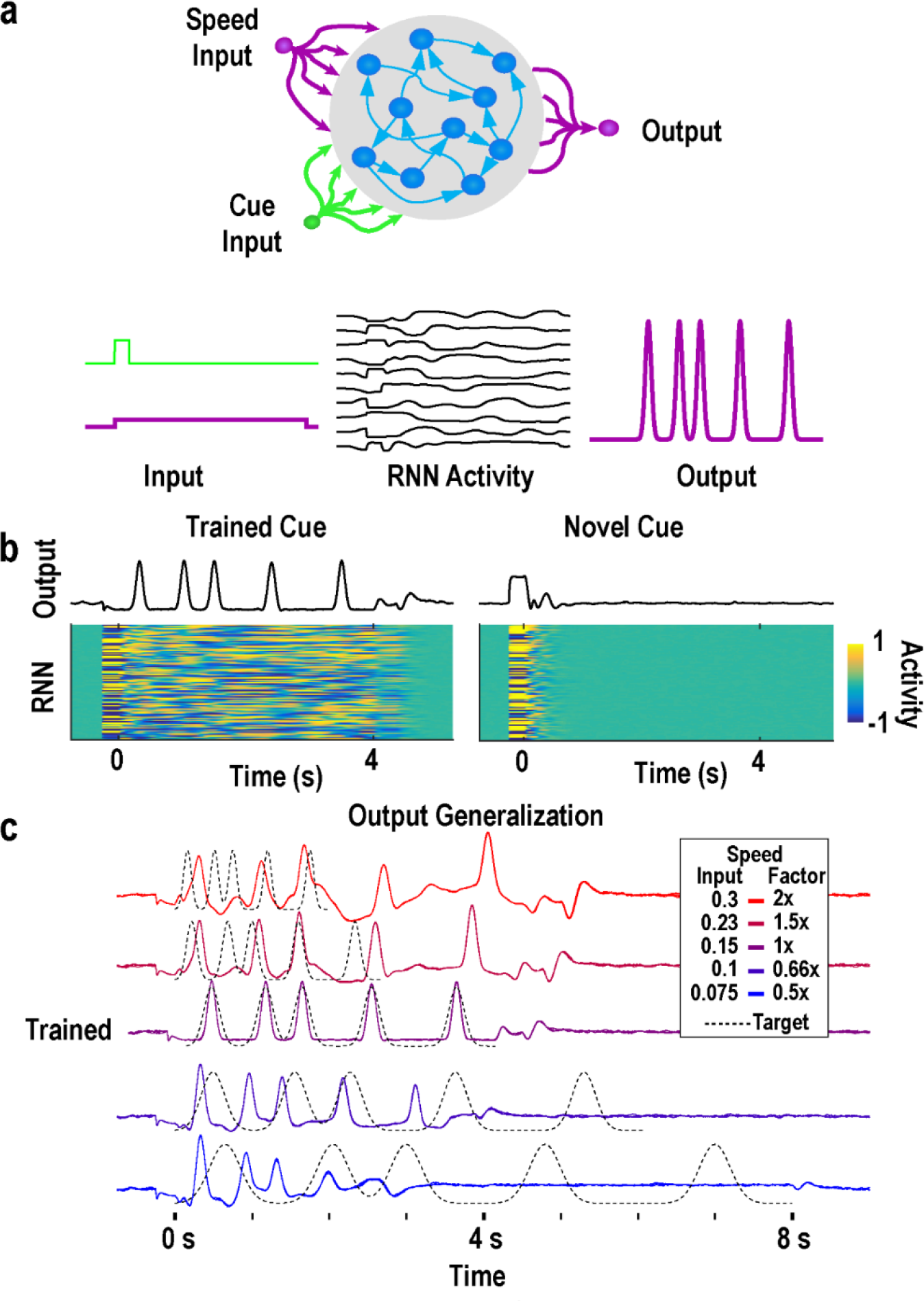
Temporal scaling is not produced by altered input drive of a RNN model. **a)** The model was composed of sparse recurrently connected firing rate units, which received input from two independent inputs and connected to a single output. One input served as a start cue and was active briefly at the start of each trial between *t* = [-250,0] *ms*. The second input delivered a constant low amplitude speed signal for the duration of a trial. **b)** The recurrent networks were trained to autonomously produce a neural trajectory lasting four seconds at 1x speed (*In_SI_* = 0.15). At the end of the trajectory, the recurrent network was trained return to a “rest state” (*r* = 0), forming a “gated” attractor: networks only generate long-lasting stable dynamic activity in response to the trained cue. Following recurrent training, the output unit was trained to produce a series of five “taps” at 325, 1025, 1500, 2400, and 3500 ms. In response to a novel cue input the RNN activity does not enter the trained dynamic attractor, and activity quickly returns to rest. *c)* Networks trained at one speed do not scale the speed of their dynamics in response to changing input drive. The speed signal was varied between *In_SI_* = [0.3,0.23,0.15,0.1,0.075] corresponding to 2x, 1.5x, 1x, 0.66x, and 0.5x speeds. Dotted lines illustrate the expected output pattern with correct scaling. The network speed accurately reproduces the output pattern only at the trained speed, degrading the further the speed input is from the trained value.

We trained the RNN to reproduce an “innate” pattern of network activity while receiving a speed input at a fixed amplitude (defined as speed 1x, *y_SI_* = 0.15), then trained the output to produce an aperiodic pattern composed of five “taps” after the cue onset (Online Methods). Unlike biological motor systems, RNNs in high-gain regimes are typically spontaneously active, i.e., they exhibit self-perpetuating patterns of activity. To increase the model’s congruence with cortical dynamics and motor behavior, we developed a method of training the recurrent units to enter a rest state when not engaged in a cued task. In this procedure, the recurrent units are trained to maintain a firing rate of approximately zero after the target pattern has terminated (Online Methods). This training produces a “gated dynamic attractor”: in response to a cued input the network produces the trained dynamics and then returns to a rest state (Fig. 2B). In contrast, in response to an untrained input the network activity quickly decays to the rest state. Consistent with the lack of spontaneous activity the real eigenvalues of the trained weights are less than one (Supplementary Fig. 1).

After training, the network was able to reproduce the target output at the trained speed. However, when tested at novel speeds—by changing the amplitude of the tonic speed input—the network did not exhibit temporal scaling (Fig. 2C). Notably, the network’s ability to produce the trained pattern progressively degraded with more extreme speed inputs. These findings establish that simply changing the amplitude of a tonic input does not produce temporal scaling.

### RNN Model of Temporal Scaling Predicts a Weber x Speed Effect

The above result suggests that temporal scaling is not an intrinsic property of timing in RNNs, so we next asked if temporal scaling could be learned. We trained RNNs to produce the same pattern of activity in the recurrent units at two different speeds (0.5x and 2x, Online Methods). After the recurrent network was trained, we trained the output to produce the same pattern as in Fig. 2 but only at the 2x speed. When tested at different speed input levels, the network now exhibited robust temporal scaling (Fig. 3A-B). Note that because the output was trained only at 2x speed, any change in the speed of the output reflects an underlying change in the speed of recurrent activity. RNNs trained on two speeds accurately compressed or dilated the motor pattern between the trained 2x and 0.5x speeds—in contrast to networks trained at only one speed (Fig. 3C). In other words, the network accurately interpolated to untrained speed inputs in between the trained speeds. We also tested extrapolation by testing with the speed inputs outside the trained range. Results revealed extrapolation progressively degraded with speed inputs outside the training range (Supplementary Fig. 2). These results are consistent with our results from Fig. 1: that temporal scaling is not an intrinsic property of motor timing but can be learned.

**Figure 3.**
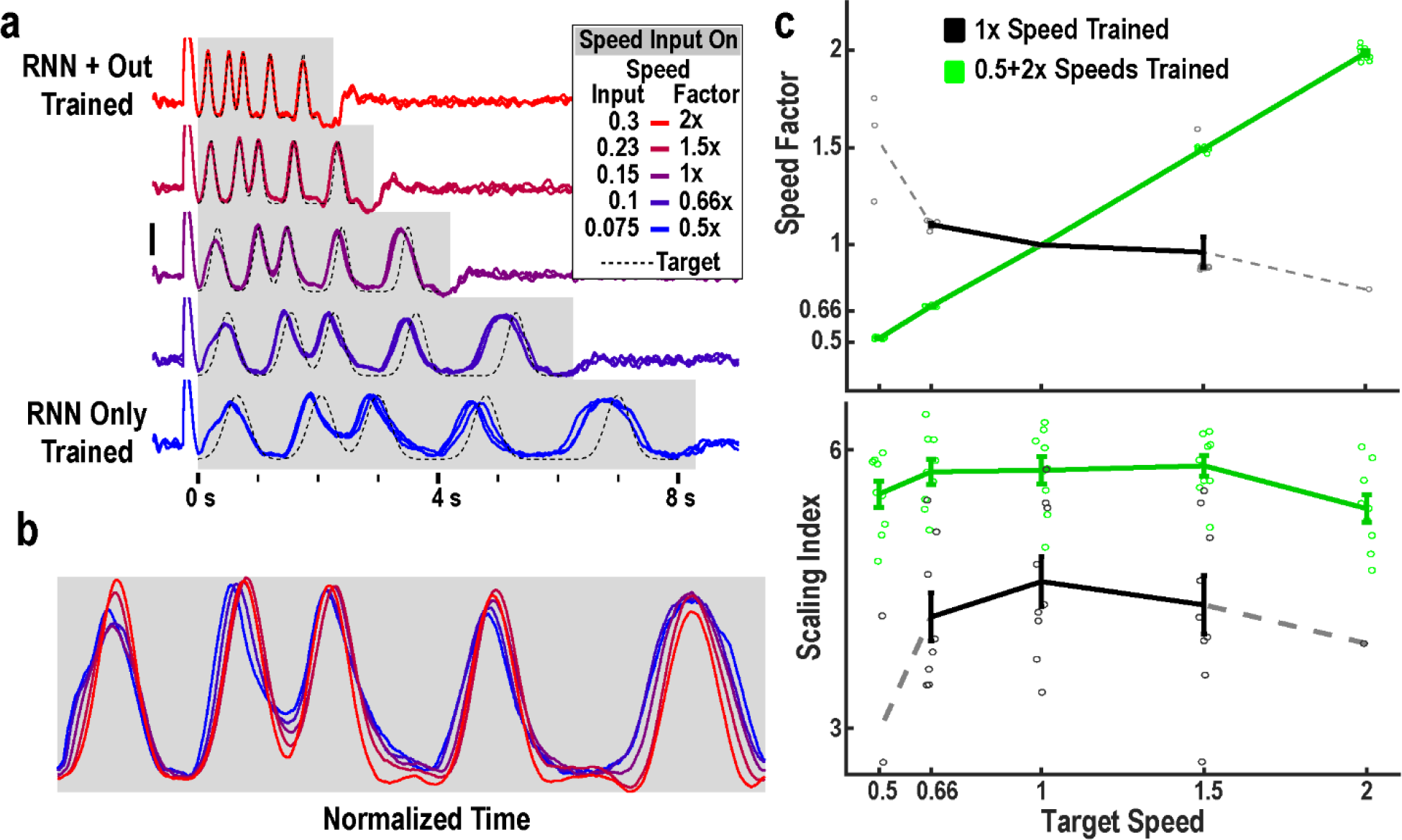
RNNs trained at multiple speeds exhibit robust temporal scaling. **a)** Output activity of an RNN trained to produce the scaled patterns of recurrent activity at 0.5x (*In_SI_* = 0.075) and 2x (*In_SI_* = 0.3) speeds. The output was trained only at the 2x speed. After training (weight modifications stopped), the network was tested at different input speed levels (*In_SI_* = [0.075,0.1,0.15,0.23,0.3])—corresponding to speeds of 0.5, 0.66, 1, 1.5, and 2x. Three example test trials at each speed are overlaid. **b)** One trial from each test speed above shown with time normalized to the end of the active period. **c)** Networks (n=10) trained at two speeds generalize to untrained speed inputs. Top: The speed factor (the mean ratio of the final tap at each speed to the mean final tap time at 1x speed over twenty trials) of networks trained at two speeds matches the target speed (green), but the speed factor of networks trained at one speed does not (black). Bottom: The scaling index (Fisher transformed correlation of the tap and target times) networks trained on two speeds is higher than those trained on one speed. Error bars represent SEMs, and circles show the value for each network. Because the activity of the one-speed networks degrades at more extreme speeds as shown in Fig. 1, many networks did not produce detectable “taps” (output peaks) at extreme speeds and we therefore could not calculate a scaling factor or index for them. We show in grey the calculations for the networks that completed at least one trial at the extreme speeds.

As mentioned above a signature of motor timing is that the standard deviation (SD) of timed responses increases linearly with time^18,22^. Because Weber’s law is often held as a benchmark for timing models^17^, we examined whether the SD of the model’s cross-trial tap times was linearly related to absolute time (the mean time of each tap in the sequence). There was a strong linear relationship between standard deviation and time (Fig. 4A) which allowed us to calculate the Weber coefficient (defined as the slope of the variance versus time squared fit). In contrast to a number of other models of timing—drift-diffusion models for example^23^—RNN based models can inherently account for Weber’s law. This in part is because the recurrent nature of these networks can amplify noise and impose long-lasting temporal noise correlations leading to near linear relationships between SD and time^24^ (Supplementary Fig. 3).

**Figure 4.**
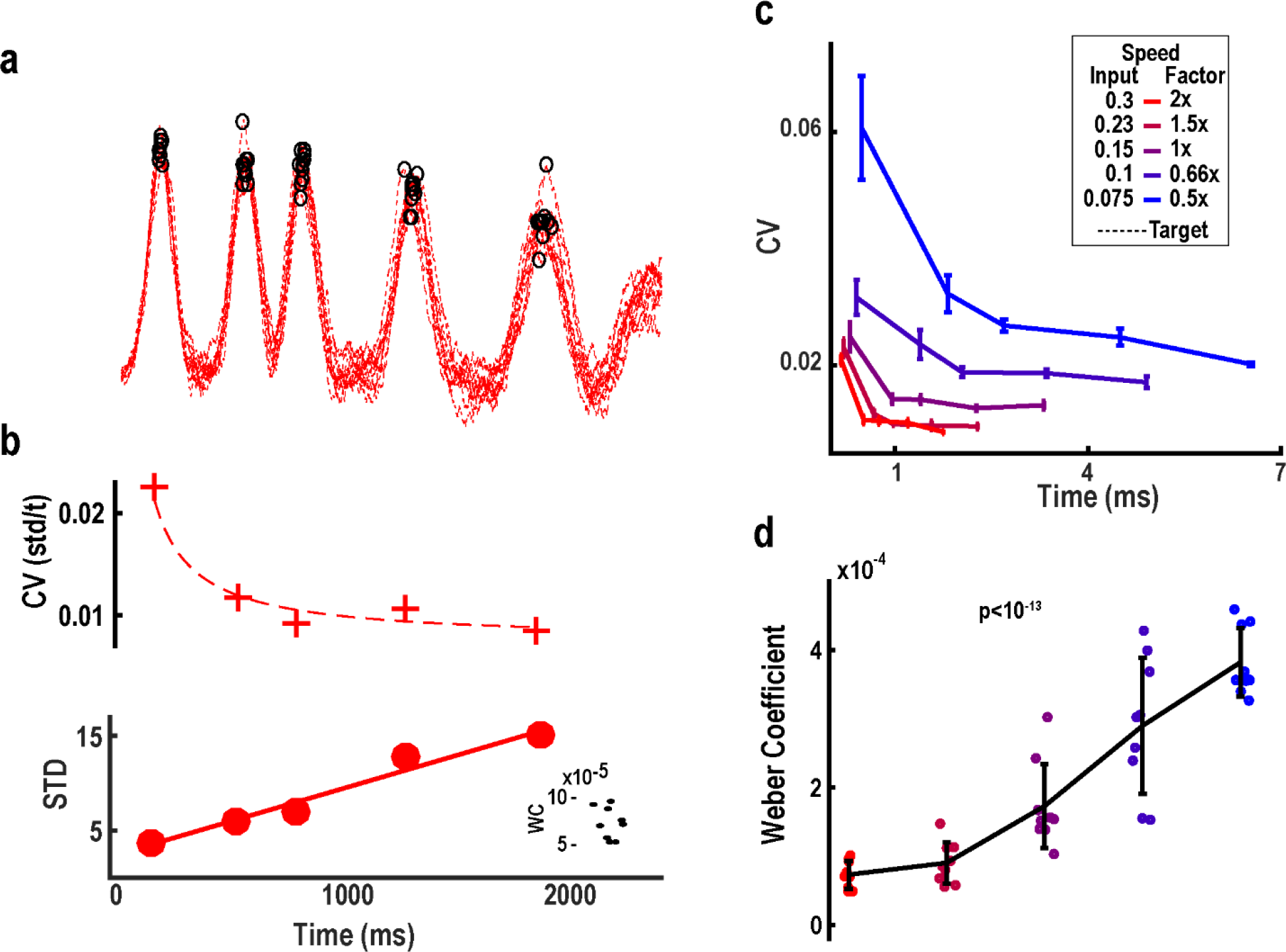
RNN models of temporal scaling predict a novel Weber-speed effect. **a)** Ten trials of the output activity of one network at 0.5x speed with tap times indicated by black circles. b) T rained RNNs account for generalized Weber’s Law, which predicts a linear relationship between the mean and standard deviation of timed intervals. Top: The coefficient of variation (CV, SD/t) at each of the five taps shown in (a). The dotted line shows the CV calculated using the fit below. Bottom: standard deviation linearly increases with time. Line shows the linear fit (r^2^=0.96). Inset shows the Weber Coefficient (the slope of variance vs. mean time) at 0.5x speed for all ten trained networks. c) The CV of ten networks calculated from twenty trials at each tested speed. Note that at the same absolute time across speeds, the CV is higher when speed is slower (the Weber-Speed effect). d) The Weber Coefficient increases at slower speeds (Repeated-measures one-way analysis of variance; F=54.4, p<10^−13^). Networks (n=10) for this analysis were trained and tested at 0.25 noise amplitude.

Across speeds there was a clear relationship between the speed and coefficient of variation (CV or the Weber fraction, Fig. 4B), as well as with the Weber coefficient (Fig. 4C)—defined as the slope of the linear fit of the variance versus *t*^2^. Specifically, the lower the speed the higher the Weber coefficient. This is a counterintuitive observation, as it implies that at the same absolute time, temporal precision is significantly worse at slower speeds. To use a clock analogy: it would be as if the same clock was more precise at timing a 2 s interval when that interval was part of a short (high speed) pattern compared to a 2 s interval that was part of a long (slow) pattern. In other words, the model predicts that humans will be less precise tapping halfway through a four second pattern, compared to the last tap produced at the end of the same pattern produced over two seconds.

### Test of Prediction: Humans Exhibit the Weber x Speed Effect

To the best of our knowledge the notion that the effects of noise on temporal precision would be more severe at low speeds (longer durations) has never been predicted or experimentally tested. Thus, we used a temporal production task in which subjects were required to reproduce an aperiodic pattern composed of six taps at five different speeds. The pattern and speeds were the same as those in the model above, with the first tone in this task acting as a self-initiated cue. Subjects (n=25) listened to an auditory pattern composed of six tones, and were asked to reproduce it using a keypad (Fig. 5A, Online Methods). In each block subjects heard the pattern at one of five temporally scaled speeds (0.5x, 0.66x, 1x, 1.5x, and 2x) and reproduced the pattern (Fig. 5B, single subject). Based on the mean and SD of the taps it is possible to calculate the CV for each tap at each speed, as well as the Weber coefficient (inset Fig. 5B, SD versus *t* is shown for visualization purposes). Under Weber’s law the CV of timed responses should be the same for taps at the same absolute time, and the Weber coefficients should be the same across speeds. However, across subjects (Fig. 5C) CVs were significantly different across speeds (F4,96=8.40, p<10^−5^, speed effect of a three-way repeated ANOVA), and the Weber coefficient decreased with higher speed (F4,96=4.18, p<0.005, one-way repeated ANOVA).

**Figure 5.**
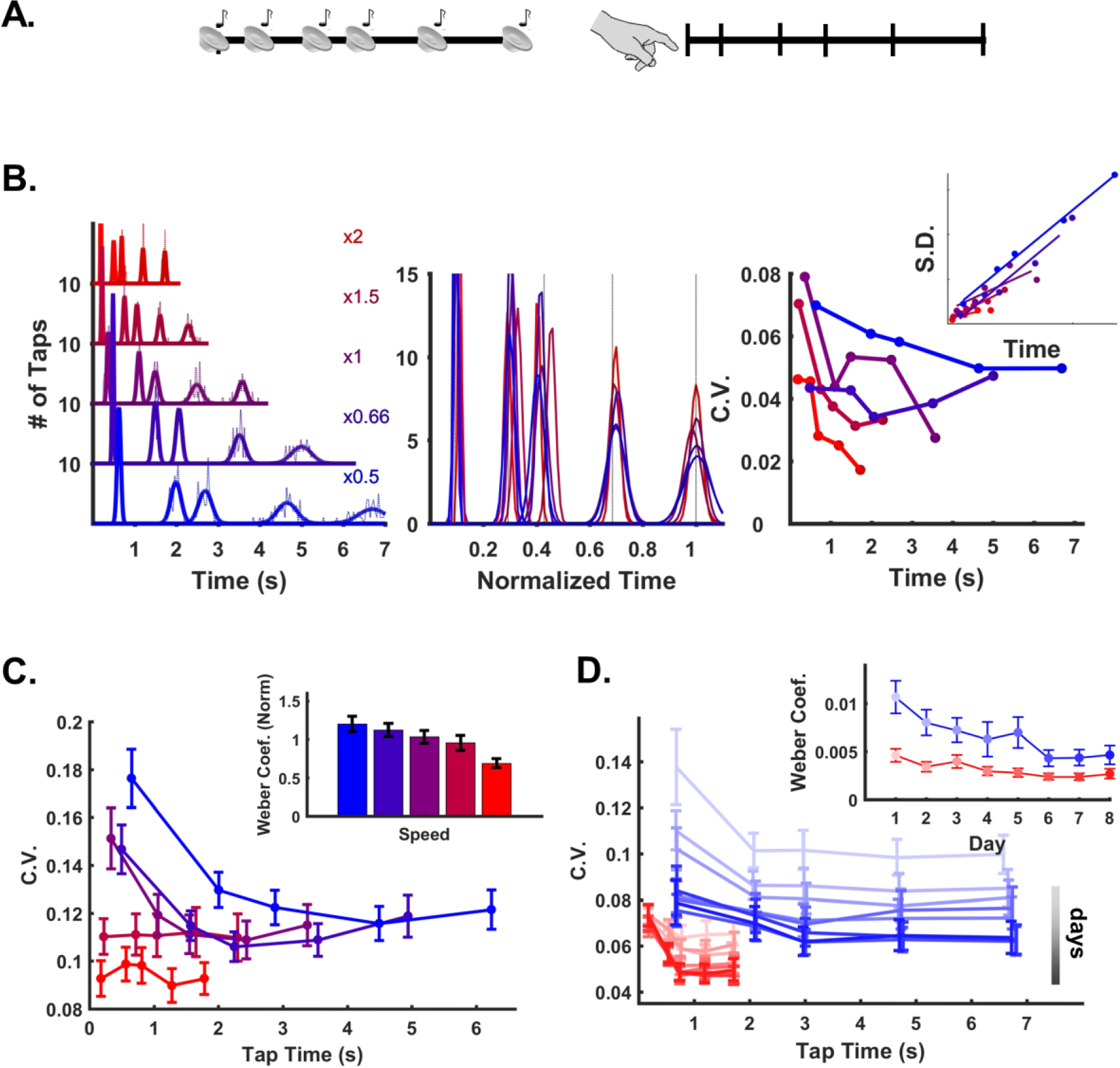
Test of the Weber-Speed effect prediction. **a)** Subjects were trained on an auditory temporal pattern reproduction task, using the same aperiodic pattern and same five speeds used to train the RNNs. **b)** Left: Histogram (dashed lines) and Gaussian fits (solid lines) of the cued taps at all five speeds from a single subject. Middle: the fits shown with time normalized to the mean of the last tap—note that the scaled fits do not overlap as expected by Weber’s law. Right: C.V. of each tap at each speed, with SD vs mean time inset. **c)** Mean and SEM of all subjects (n=25) for the five tested speeds. Note that, as in the RNN model, the CV at the same absolute time is higher at slower speeds. Inset shows the Weber coefficient at each speed. **d)** The Weber-Speed effect is not due to inexperience with the task. A subset of fourteen subjects were trained to produce the fast and slow speeds over eight additional days. The Weber-Speed effect persists over the course of training. Figure S5 plots the same data while incorporating the within group distributions.

The above data is potentially confounded with task difficulty or learning—that is, the difference in the Weber coefficients across speeds could potentially reflect some nonspecific effect in which slower patterns are harder to memorize and reproduce. We thus trained a subset of subjects (n=14) on the fastest and slowest speeds over an 8-day period. Again at the same absolute time the CV was lower for the faster speed across training days (e.g., ≈0.7 s in Fig. 5D). The Weber coefficient was significantly smaller for the faster speeds across training days (Fig. 5D, inset; F1,13=16.58, p<0.002, speed effect two-way repeated ANOVA; pairwise posthoc test on each day, maximum p = 0.056, Tukey-Kramer)—even as asymptotic learning was evidenced by progressive decrease in the Weber coefficients of both speeds across days. These results establish a novel psychophysical phenomenon, a Weber-speed effect, in which in absolute time, motor timing is more precise when executed at faster speeds.

### Mechanisms of Temporal Scaling

Having established a model of temporal scaling that generated a correct prediction we next used the model to examine the potential network-level mechanisms underlying temporal scaling. At first glance the notion that an RNN can generate the same trajectory at different speeds is surprising, because it seems to imply that different tonic input speed levels can guide activity through the same points in neural phase space at different speeds. Furthermore, it is important to emphasize that the model exhibits temporal scaling whether the network is trained so that larger speed inputs increase *or* decrease trajectory speed. We first visualized the trajectories in PCA space, which revealed that the trajectories at different speeds are not overlapping, but follow a slightly offset path through neural phase space (Fig. 6A). In other words, the trajectories are arranged according to speed in an apparently “parallel” manner. To quantify this observation we calculated the Euclidean distance in neural space (N=1800) between the trajectory at each speed and the 0.5x speed (Fig. 6B). Finding the minimum distance between the comparison speed and the 0.5x speed (the “reference” trajectory) revealed that the trajectories maintain a relatively constant distance from each other (Fig. 6C). This measure provided an unbiased estimate of the relative speed of each trajectory. For example, if the test trajectory is moving four times as fast as the reference trajectory, it should be closest to the reference trajectory when it has been active for ¼ the elapsed time. In other words, plotting *t*^2*x*^vs 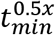should form a line with slope 0.25, which is indeed what we observed. Moreover, this relationship generalized to novel interpolated speed levels (Fig. 6D).

**Figure 6.**
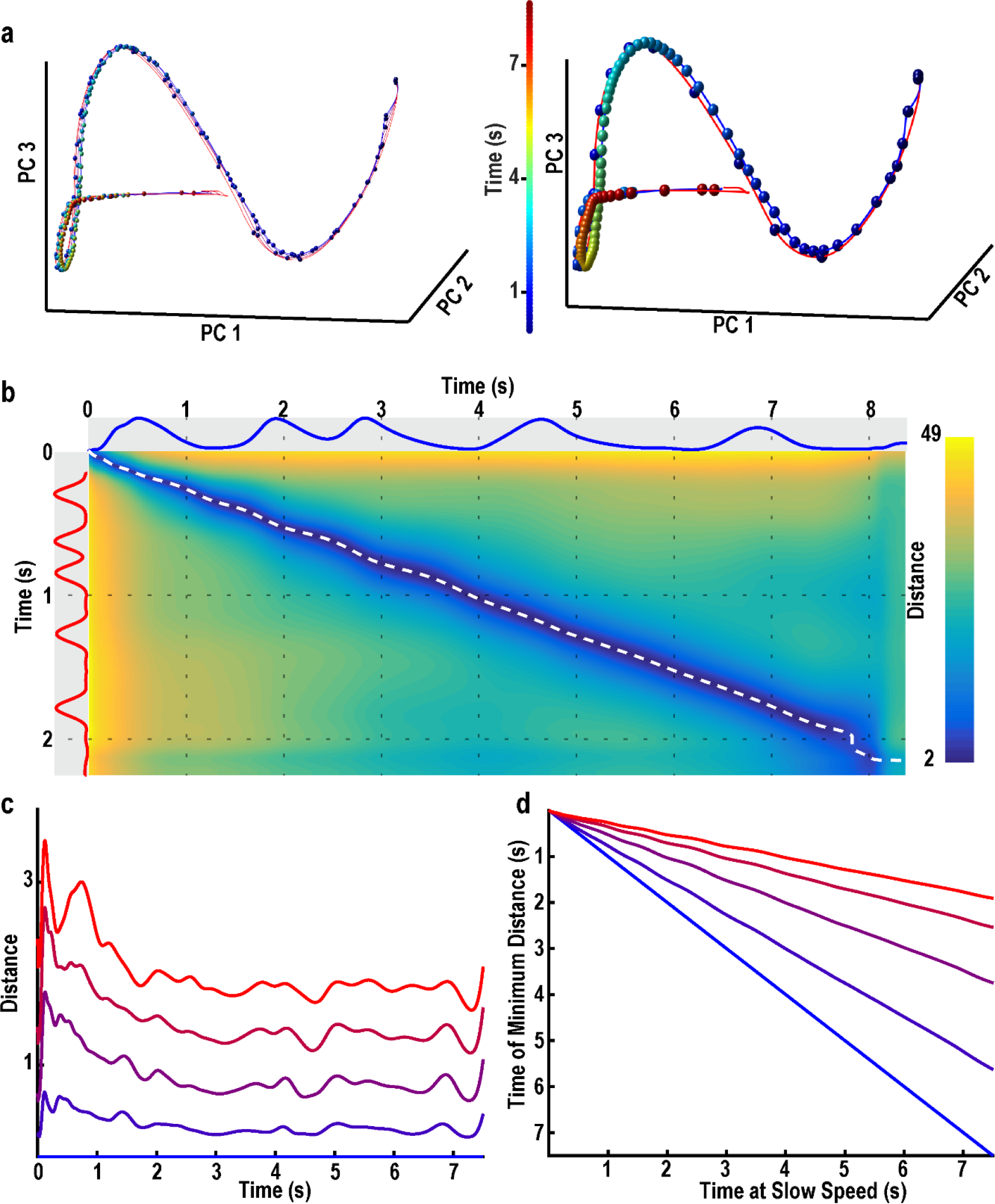
Temporal scaling relies on parallel neural trajectories at different speeds. **a)** Trajectory of RNN activity at five speeds projected onto the first three principal components. Right: same data, but only the slowest (blue line) and fastest (red) speeds are plotted to highlighted the difference in speed of the two trajectories. Colored spheres indicate absolute time in each trajectory, with 100 ms between spheres.**b)** Euclidean distance matrix between the fast and slow trajectories in neural space at each point in time along each trajectory (network size: N=1800). Blue and red traces along the axes show the output. White dotted line traces the minimum distance between the two trajectories, which never reaches zero.**c)** The minimum distance along the slowest trajectory from each other speed. **d)** The relative timing at which the minimum occurs in each trajectory. For example, the slowest, 2x speed and 1x speed trajectories are closest after 4 s of the 2x trajectory and 2 s of the 1x trajectory.

The above finding presents an interesting mechanism of how trajectories of different speeds are formed in state space, but does not address how the magnitude of a static speed signal can temporally scale the time-varying dynamics of a nonlinear RNN. Understanding the underlying dynamics of complex nonlinear neural networks is a notoriously challenging problem with only a few tools available^25^. Here we introduce a method to dissect the internal forces driving a network. We first quantified the total drive to the network: the time-dependent change in the total synaptic input onto each neuron in the RNN. Measuring the magnitude (Euclidean norm) of the total drive showed that— in contrast to untrained networks or to networks trained at a single speed—the total drive scaled with the cued speed (Fig. 7A). To address how the total drive scales the neural dynamics, we used a novel network drive decomposition method^26^. This approach decomposes the total network drive into its three components: the recurrent synaptic drive, the synaptic decay (which is always driving the network towards the origin), and the external tonic (time-independent) speed input (see Fig. 7C). While the speed input magnitude scales with speed as defined by the experimental conditions, neither the recurrent nor the decay magnitudes scaled with speed, meaning that the recurrent or decay components in isolation cannot account for temporal scaling (Fig. 7B).

**Figure 7.**
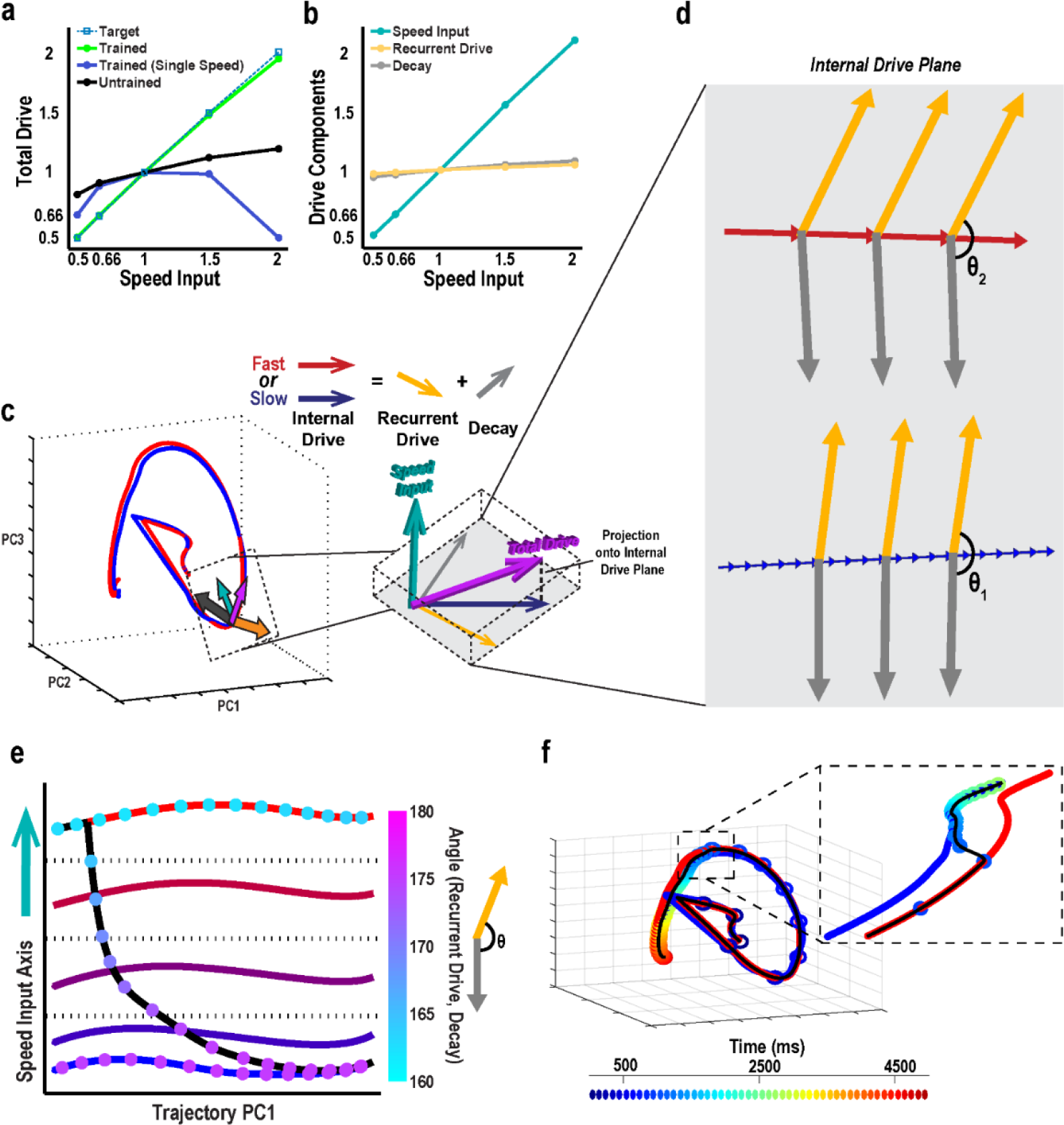
Mechanisms of temporal scaling in the RNN. **a)** Magnitude of the instantaneous change in activity of the recurrent network (total drive) scales linearly with speed in networks trained at two speeds, but not in networks trained at one speed or untrained networks. Total drive is normalized to the 1x speed. **b)** Decomposing network drive into its three components (recurrent, decay, and input) reveals that the recurrent and decay components do not individually scale with speed, thus neither of them in isolation can account for temporal scaling. **c)** To examine the relationship between the recurrent and decay components separate from the input drive, we projected them onto the internal drive plane, a subspace orthogonal to the speed input (Online Methods). **d)** This projection revealed that at faster speeds the angle between the recurrent and decay components decreases, creating a second-order effect that drives the network activity along the trajectory more quickly. **e**) Network activity projected onto the input axis and the first principal component of network activity (the dimension which accounts for the largest amount of variance). The colored markers indicate the angle between the recurrent and decay components. The position along the input axis does not change as a function of time, indicating that speed is encoded by the position along the input axis. When the speed input level is abruptly decreased partway through the trajectory (black line), the network switches from fast to slow speeds via an increase in the angle between the recurrent and decay components. **f)** Neural trajectories in the first three principal components during a mid-trajectory change in speeds. As the dynamics transition from fast to slow (inset), the trajectory moves along a hyperplane defined by the parallel trajectories shown in Fig. 6.

Analysis of the dynamics also revealed that, at each speed, the trajectory traverses directions that are independent of the speed input—i.e., the projection of each trajectory onto the speed input axis exhibits small variance (explained variance was <1% at all speeds, Fig. 7E). There are two consequences to the observations that the time-varying dynamics are not driven by the input, and that the recurrent drive and decay magnitudes do not exhibit temporal scaling: 1) at each speed, some combination of these internal drive components counterbalance the speed input; and 2) they *collectively* underlie temporal scaling of the trajectory. In order to isolate the contribution of these interactions we studied the internal drive components in the subspace orthogonal to the speed input axis (Fig. 7C). Measurements showed that even in this subspace, changes of recurrent drive and decay magnitudes do not explain temporal scaling of the total drive (data not shown). Instead, the recurrent synaptic drive and decay oppose each other (the angle between them is obtuse) throughout the trajectory, and the extent of this opposition alters the trajectory’s speed (Fig. 7D). Specifically, the angle between the two components decreases as the speed input increases (θ_2_ < θ_1_), thereby amplifying the net (or total) drive.

Projecting the trajectories onto the speed input axis revealed that speed is encoded in the trajectory’s position rather than its direction (Fig. 7E). Moreover, by traversing phase space along directions that are independent of the speed input, the trajectory’s position with respect to the speed input stays relatively constant. This allows the speed of the trajectories to stay relatively constant. To confirm this, we asked if—as with biological motor patterns—a network could switch speeds mid-trajectory. Indeed, by decreasing the speed input in the middle of a fast (2x) trajectory we observed a rapid transition to the slow trajectory (Fig. 7F). Network drive decomposition showed that a change in the speed input causes an imbalance between it and the internal drive, altering the position of the trajectory along the speed input axis. In turn, this increases the angle between the recurrent drive and decay, which slows the trajectory down. It also rebalances the speed input and the internal drive components such that trajectory speed stops changing when the input to internal drive balance is restored (Fig. 7E). Together, these results demonstrate that temporal scaling is the outcome of speed input-dependent balance between the recurrent and decay drives.

## DISCUSSION

It is increasingly clear that on the scale of hundreds of milliseconds to seconds the brain represents time as dynamically changing patterns of neural activity ^4,5,12,14,27^. Timing on this scale exhibits a number of properties including: 1) the ability to execute the same motor patterns at different speeds, and 2) temporal variability increases linearly with time (Weber’s law). Here we unify and extend these observations by presenting a recurrent neural network model that not only performs temporal scaling and accounts for Weber’s law, but also predicts that Weber’s law is speed-dependent. Specifically, the Weber coefficient—the slope of relationship between SD and absolute time—is speed-dependent. We tested this prediction using human psychophysics experiments, and confirmed that in absolute time the temporal precision of motor response is dependent on speed.

### Weber’s Law

While Weber’s law is a well-established property of timing in humans ^28–30^ the neural underpinnings of Weber’s law have long been debated ^23^. Early models of timing (internal clock models) consisted of an accumulator that integrated the number of pulses of a noisy oscillator. In their simplest form, however, these models did not account for Weber’s law because the standard deviation of such a clock will increase as a function of 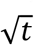 rather than *t.* Thus internal clock models postulated that Weber’s law arises from a second clock-independent noise source, such as the memory of the interval being generated^18,31^. Other models, like those based on the variance between multiple timers, can intrinsically account for Weber’s law ^23,32^, but the biological plausibility of such variance-based models remains an open question. Our results suggest that population clock models based on recurrent dynamics can also intrinsically account for Weber’s law. This is likely due to their recurrent nature, which actively amplifies noise through internal feedback.

As other groups have pointed out, this raises the question: if independent noise sources cause temporal precision to decrease as a function of 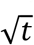, why does the nervous system settle for Weber’s law^24^? First, it is possible that this reduced accuracy is an unavoidable consequence of the abundance of correlated noise on time scales at which the intervals being timed^33^. For example, in any neural circuit, slow fluctuations produced by sensory inputs or other brain areas will impose local temporal correlations. Second, the very nature of recurrent neural networks can amplify noise by internal feedback, making Weber’s law a necessary cost of the increased computational capacity provided by such architectures.

### Weber-Speed Effect

The strong relationship between motor speed and the Weber coefficient raises the question of why—to the best of our knowledge—a Weber-speed effect has not previously been reported. One reason is that the vast majority of timing studies have relied on interval or duration tasks rather than pattern timing. Thus, in most studies the Weber coefficient is calculated by fitting the variance of timed responses of distinct intervals collected across blocks. With this approach it is not possible to explicitly address questions pertaining to temporal scaling or observe a differential effect of speed on the Weber coefficient. In contrast, by looking at the timing of complex motor patterns consisting of multiple taps^34^, it is possible to estimate the Weber coefficient within each condition (speed), revealing a strong dependence of the Weber coefficient on speed. A potentially related phenomenon bears noting: that subdivision, e.g. tapping your foot to a beat, can improve timing accuracy^31,35^. The typical interpretation of this effect is that it results from the coupling of Weber’s law and a reset occurring at each subdivided interval. Such reset mechanisms, however, result in a sublinear relationship between SD and time in the task studied here^34^. We thus hypothesize that subdivision effects may in part reflect the speed of the underlying neural trajectories.

As with Weber’s law, the Weber-speed effect raises the question of why the nervous system would utilize a timing mechanism that is inherently better—more precise across trials—when engaged in a fast motor pattern versus a slow pattern. Again, the answer lies in part in the inherent properties of recurrent circuits. An analysis of the temporal noise correlation revealed larger and longer lasting noise covariance in the RRN during slow trajectories compared to fast trajectories (Supplementary Fig. 4). Additionally, there is an inherent inverse relationship between the speed of a dynamical system and the effects of noise. Consider a sinusoidal function at a high (short period) and low (long period) speed in the presence of additive noise. If we were to consider each peak in the amplitude of the function as a tic of a clock, additive noise will produce more variance in the peaks of the slow curve because noise added to a slowly changing function is more likely to change the times of the peaks.

### Predictions

The model of temporal scaling presented here makes a number of experimental predictions. The most important prediction, that movements executed at higher speeds are more temporally precise in absolute time, has been tested and confirmed. However, a number of important questions pertaining to this prediction remain, including whether simple interval production tasks correspond to executing the same neural trajectories at different speeds. At least one study in the striatum suggests that during a timing task it is indeed the case that different intervals are timed by the same neural patterns unfolding at different speeds^4^. However, no electrophysiological studies have examined temporal scaling during the production of aperiodic temporal patterns similar to those studied here.

The model makes a number of additional neurophysiological predictions. First, it should be possible to use electrophysiological recordings during temporally scaled motor behavior to observe neural trajectories whose positions on a manifold (a “surface”) in high-dimensional space reflect the speed of the motor pattern. Second, the model predicts longer temporal correlations in slower trajectories, and thus that slower trajectories should exhibit longer lasting temporal noise covariance. In other words, on a trial-by-trial basis, when the population clock reads early at the beginning of a trajectory, that deviation will persist longer if the trajectory is moving slowly.

While we propose that the model presented here captures general principles of how neural dynamics account for timing and temporal scaling, the learning rule used to generate the neural trajectories driving timing is not biologically plausible. Future research will have to determine whether such regimes can emerge in a self-organizing manner. Additionally, while the model is agnostic to what parts of the brain generate such patterns, we hypothesize that similar regimes exist in neocortical circuits characterized by recurrent excitation. However, this model does not exclude that brain circuits defined by recurrent inhibition—the cerebellum^6,36^ and striatum^37,38^—may also be responsible for temporal scaling. Indeed, a recent model of timing in the striatum has proposed how recurrent inhibition could account for temporal scaling^39^.

Overall the current studies support the notion that many neural computations can be implemented not by converging to a point attractor^40,41^, but as the voyage through neural phase space^42–44^. And more specifically, that these trajectories represent dynamic attractors, which can encode motor movements and are robust to perturbation—that is, they can return to the trajectory after being bumped off^8^. Here we show that recurrent neural networks can exhibit regimes with parallel families of neural trajectories that are close enough to drive the same motor pattern but that are traversed at different speeds— accounting for temporal scaling. These regimes predict that the temporal precision of motor responses in absolute time is dependent on speed of execution. This prediction was confirmed in human timing experiments, establishing a novel psychophysical Weber-speed effect.

---

**Supplementary Fig. 1.**
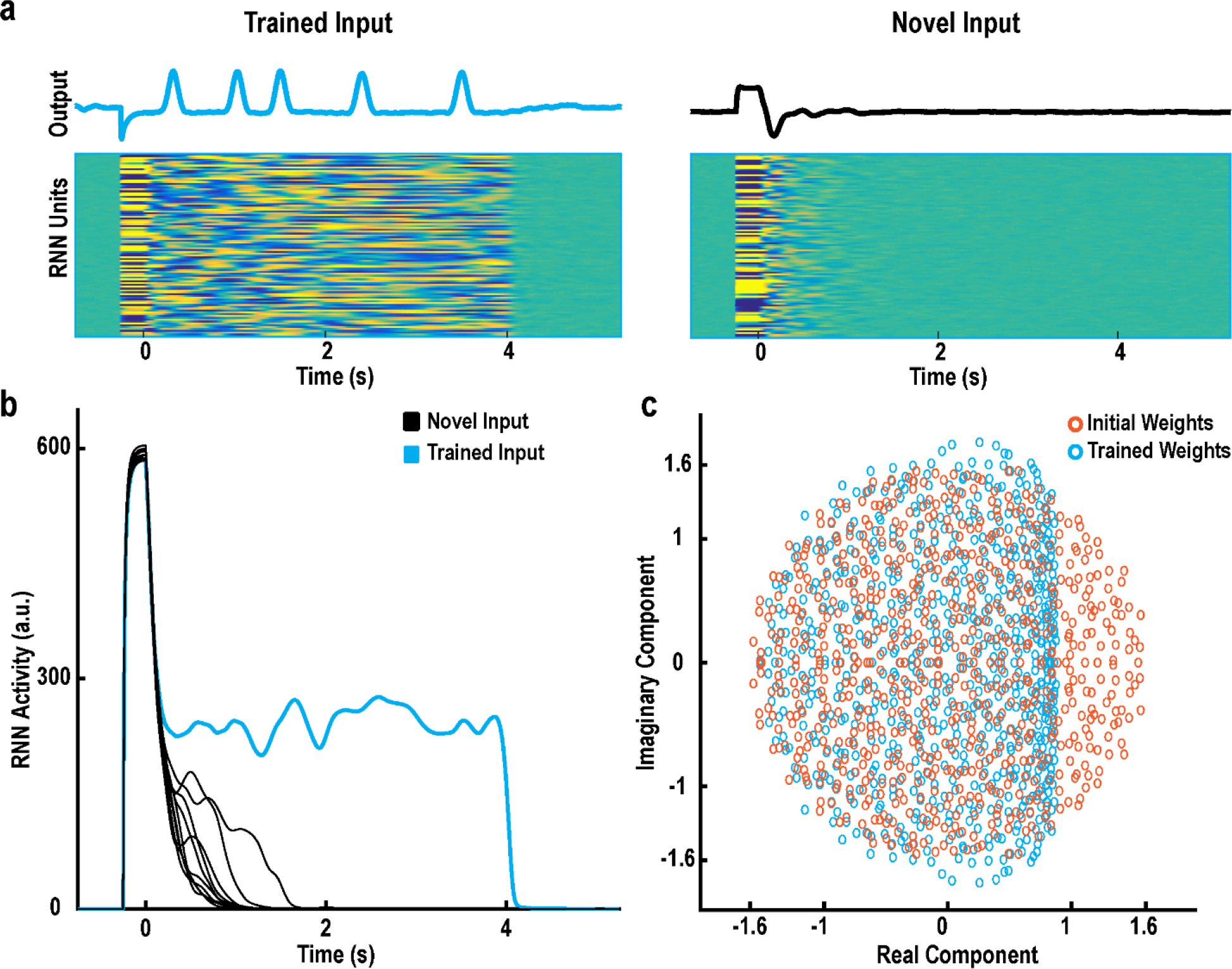
Gated attractor networks suppress untrained activity. **a)** RNNs were trained to suppress activity except in response to trained cue inputs. Left: Example activity after training in response to the trained cue input. Right: Response to an untrained cue. **b)** RNN activity (the norm of the firing rate (r) in response to the trained cue and ten untrained cue inputs. **c**) The eigenvalues of the recurrent weights before and after training. After training, the real components of the eigenvalues are less than one, meaning gated attractor networks are not spontaneously active.

**Supplementary Figure 2.**
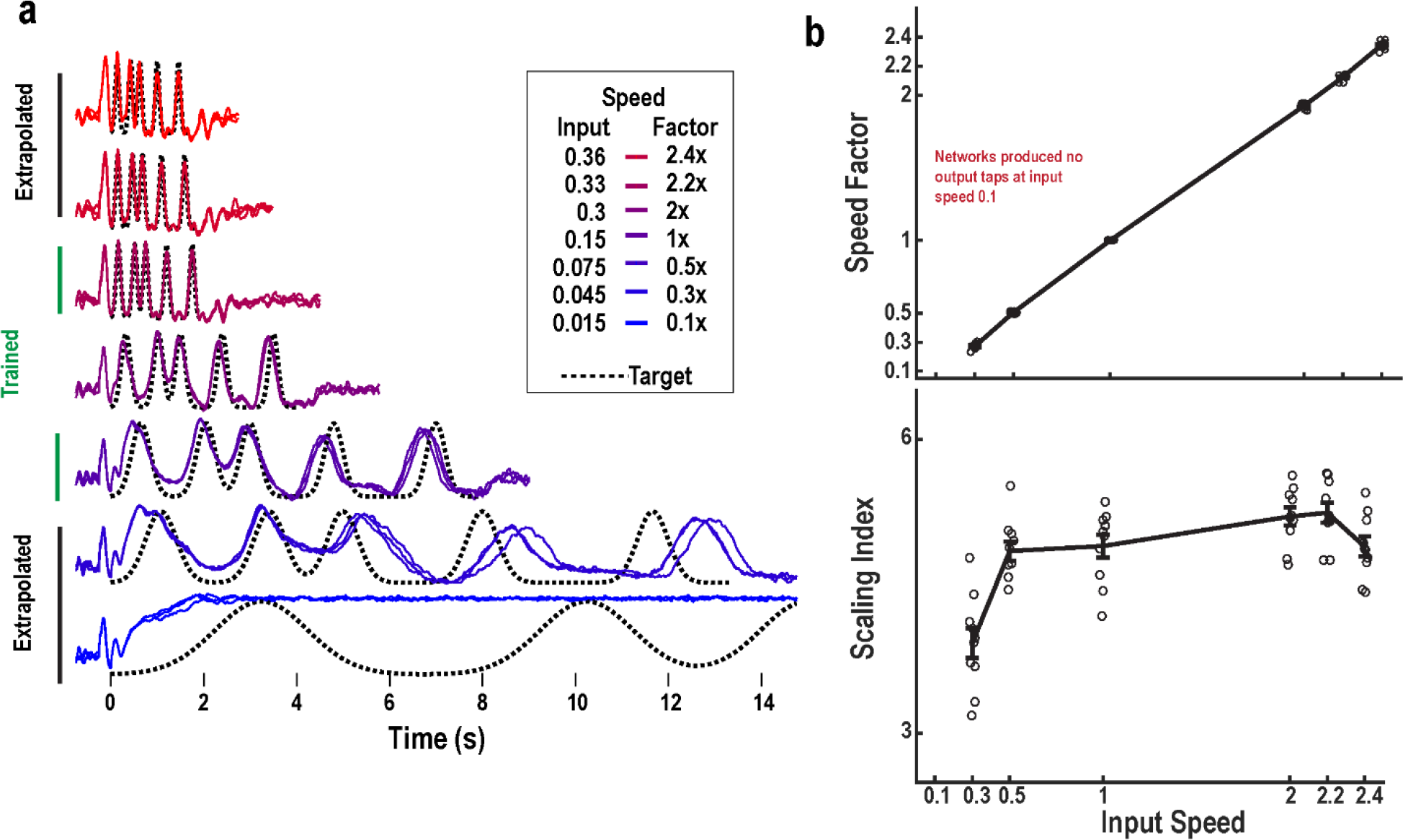
RNNs temporal scaling degrades outside of the trained speed range. **a)** Output traces at interpolated and extrapolated speeds (outside of the trained speed range). **b)** Generalization factor of ten networks calculated from twenty trials at the speeds shown above. Generalization degrades at slower speeds. **c)** The scaling factors of the ten networks.

**Supplementary Figure 3.**
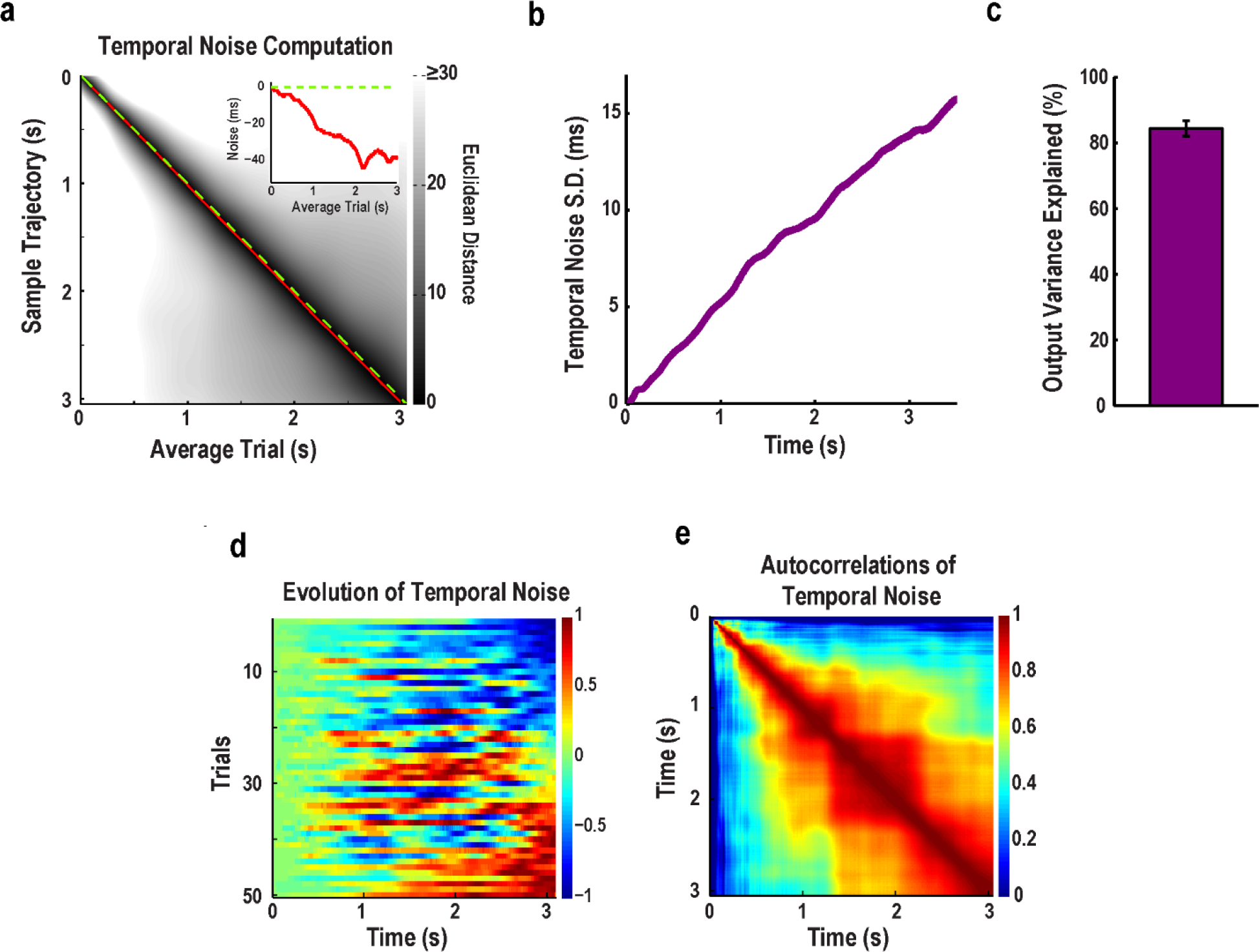
RNNs produce long-lasting temporal noise correlations. **a)** Euclidean distance matrix between the trial-averaged trajectory of a trained RNN and a single trial at speed 1x. The times at which the two trajectories are closest is represented by the red line. Matched time points between the trajectories is represented by the identity line (green dashed line). Over time the sample trajectory runs further ahead of the average trajectory (temporal noise), as evidence by the red line being below the green line. Inset: Temporal noise (red line) in the sample trajectory relative to the average trajectory. At the end of the average trajectory, the sample trajectory is ∼40 ms ahead. **b)** The linear relationship of the SD of temporal noise in the trajectories and absolute time underlies Weber’s law observed in the output unit. The SD of temporal noise is calculated over 50 trials, averaged across 10 networks. **c)** Output timing variability explained by temporal noise in the RNN (normalized mean squared error calculated between temporal noise in the trajectories and the output), averaged across 10 networks (error bars indicate SEM). **d)** Normalized temporal noise across 50 trials, sorted according to the noise at the end of the trajectory in one network. **e)** Autocorrelation of temporal noise in one network. Each element in the matrix represents the correlation (across trials) of temporal noise at the corresponding pair of time points (i.e., pair of columns in d). Deviations at early time points predict later deviations. Networks (n=10) trained and tested at noise amplitude of 0.25.

**Supplementary Figure 4.**
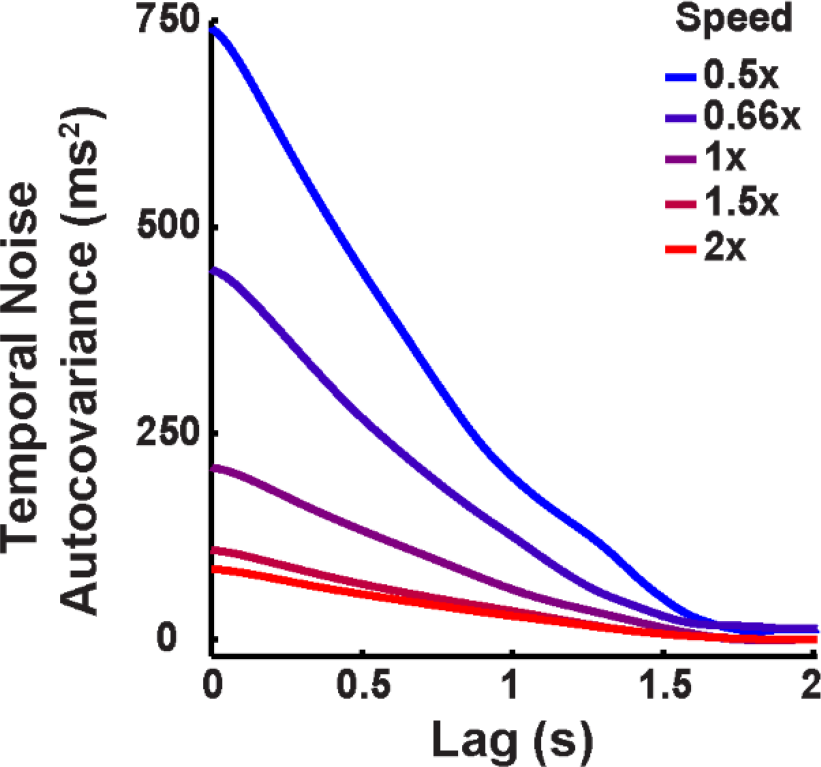
Slower speeds have longer lasting noise correlations. Autocovariance of temporal noise at different speeds across 50 trials, averaged across 10 networks; note that the slowest speed has an elevated covariance even at a time lag of up to 1 s. Networks (n=10) trained and tested at noise amplitude of 0.25.

**Supplementary Figure 5.Same data as Figure 5.**
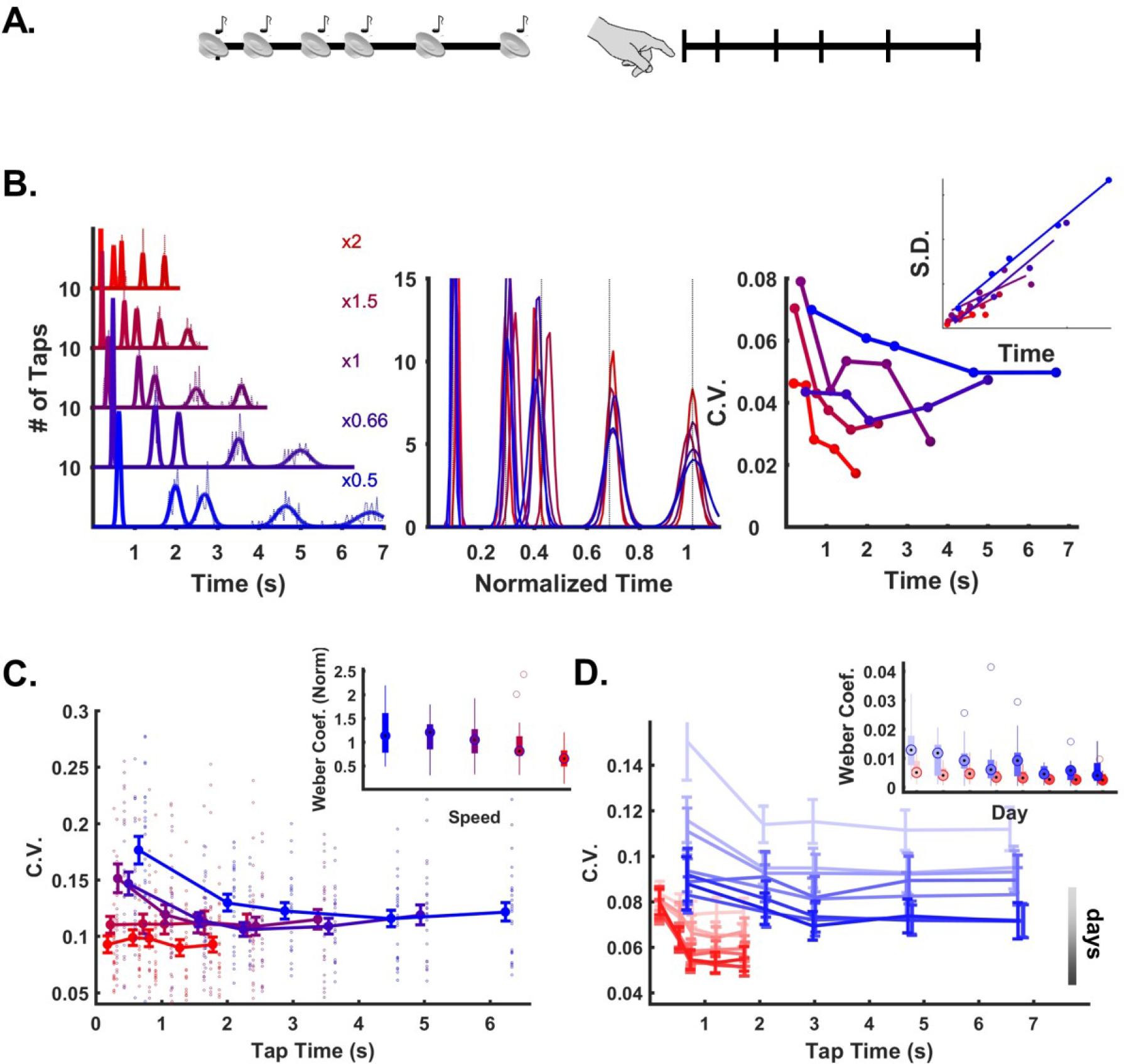
**a)** Subjects were trained on an auditory temporal pattern reproduction task, using the same aperiodic pattern and same five speeds used to train the RNNs. **b)** Left: Histogram (dashed lines) and Gaussian fits (solid lines) of the cued taps at all five speeds from a single subject. Middle: the fits shown with time normalized to the mean of the last tap—note that the scaled fits do not overlap as expected by Weber’s law. Right: C.V. of each tap at each speed, with SD vs mean time inset. **c)** Mean and SEM of all subjects (n=25) for the five tested speeds. Note that, as in the RNN model, the CV at the same absolute time is higher at slower speeds. Inset shows the Weber coefficient at each speed. **d)** The Weber-Speed effect is not due to inexperience with the task. A subset of fourteen subjects were trained to produce the fast and slow speeds over eight additional days. Boxplots (5C-D): the inner circle represents the median, box limits represent the lower and upper quartiles, the whiskers represent the 1.5x quartile range, and isolated circles are outliers.

## ONLINE METHODS

### Temporal Scaling of Motor Patterns in Humans

Human psychophysics experiments were performed using a temporal pattern reproduction task^34^. During the experiments, the subjects sat in front of a computer monitor with a keyboard in a quiet room. On each trial, subjects heard a temporal pattern and then reproduced this pattern by pressing one key on a Cedrus Response Pad^™^. The target stimulus consisted of a series of tones (800 Hz). After the subjects reproduced the pattern, a visual representation of the target and of the subject’s response appeared on the screen along with a score based on the correlation between the target and the reproduced pattern. Stimulus presentation and response acquisition were controlled by a personal computer using custom Matlab code and the Psychophysics Toolbox^45^. All experiments were run in accordance with the University of California Human Subjects Guidelines.

To test whether temporal scaling is an innate property of motor behavior, we trained subjects to produce the Morse code spelling of “time” at 10 words per minute (Fig. 1). Training occurred over four days, with 15 blocks of 15 trials per day. On the fifth day, subjects were asked to produce the trained pattern at 0.5x, 1.0x and 2x the speed under freeform conditions: Subjects first completed 15 trials of the trained pattern, and then were asked to produce the same pattern at the same speed (1x), twice as fast (2x), and twice as slow (0.5x) in the absence of any additionally auditory stimuli. Subjects performed five blocks with five trials per speed in a random order for a total of fifteen trials per block. The subjects were 10 undergraduate students from the UCLA community who were between the ages of 18 and 21. Subjects were paid for their participation.

To test the Weber-speed prediction of the RNN model (Fig. 5), subjects performed a temporal reproduction task, wherein they heard a pattern of six tones (each lasting 25 ms) and were asked to reproduce the timing of the onset of each tone with a self-initiated start time (representing the first tone). For the 1x speed the six tones were presented at 0, 325, 1025, 1500, 2400, and 3500 ms. This pattern was then scaled to five logarithmically distributed speeds: 0.5x, 0.6x, 1x, 1.5x, and 2.0x. Subjects completed four blocks of fifteen trials per speed in a random order. A randomly chosen subset of the subjects were trained to produce the 0.5x and 2x speeds over eight additional days, consisting of ten blocks of fifteen trials per speed. The subjects for this study were 25 undergraduate students from the UCLA community between the ages of 18 and 21 and paid for their participation.

### Network Equations

The units of the RNNs used here were based on a standard firing rate model defined by the equations ^46,47^:

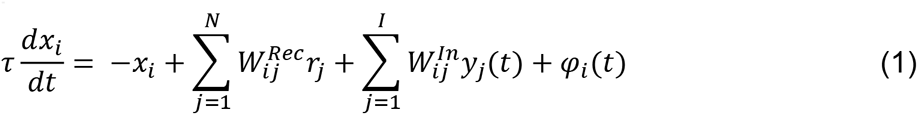

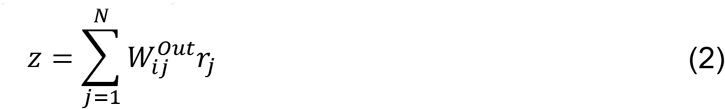

where *r_i_* = tanh(*x_i_*) represents the output, or firing rate, of recurrent unit (*i* = 1, *…,N).* The variable *y* represents the activity level of the input units, and *z* is the output. *N* = 1800 is the number of units in the recurrent network, and *τ* = 50 *ms* is the unit time constant. The connectivity of the recurrent network was determined by the sparse *N x N* matrix *W*^Rec^, which initially had nonzero weights drawn from a normal distribution with zero mean and SD 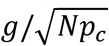. The variable *p_c_ = 0.2* determined the probability of connections between units in the recurrent network which were drawn uniformly at random, and *g* = 1.6, represents the “gain” of the recurrent network^21,48^. The *N x I* input weight matrix *W^ln^* was drawn from a normal distribution with zero mean and unit variance. For all figures, *I* = 2, except Supplementary Fig. 1, where additional input units were added to test the specificity of the network response to untrained cue inputs. One input served as cue to start a trial and its activity was set to zero except during the time window −250 ≤ *t* ≤ 0, when its activity was equal to 5.0. The second input unit served as a speed input and was set to a constant level during the time window −250 ≤ *t* ≤ *T*, where *T* represents the duration of the trial. Each unit in the recurrent network was injected with noise current *φ_i_* (*t*), drawn independently from a normal distribution with zero mean and SD 0.05, except for the Weber experiments where the SD was 0.25. The recurrent units were connected to the output unit *z* through the *Nx1* vector *W^Out^*, which was initially drawn from a normal distribution with zero mean and SD 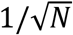.

### Recurrent Learning Rule

The networks in this study were trained using the Innate Learning Rule, which trains an initially chaotic recurrent network to autonomously yet reliably produce an arbitrary activity pattern in the presence of noise^8^. It is based on the Recursive Least Squares (RLS) update rule ^49,50^. The recurrent weights onto unit *i* were updated every *Δt* = 5 *ms* as dictated by

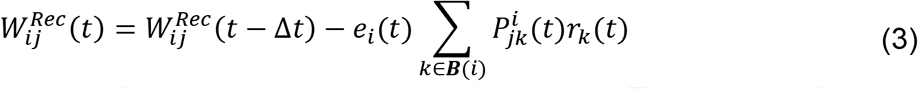

where ***B**(i)* is the subset of recurrent units presynaptic to unit *i*. The error *e_i_* of unit *i* is given by

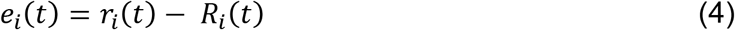

where *r_i_* is the firing rate of unit *i* before the weight update, and *R* is the target activity of that recurrent unit. The square matrix ***P**^i^* estimates the inverse correlation of the recurrent inputs onto unit *i*, updated by

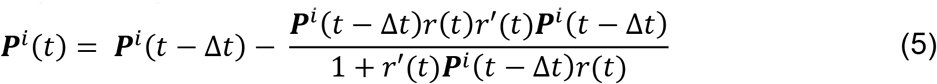

### Training Procedure

To train a network to perform the temporal scaling task, we first generated a target pattern of recurrent activity by stimulating the network with the cue input and capturing the dynamics generated according to equation (1) over 2000 ms in the presence of speed input level *y_SI_* = 0.3 and zero noise (similar results are obtained if the target pattern is harvested in the presence of noise). We then produced a temporally dilated version of this target by linearly interpolating by a factor of four to produce a second scaled version of the target with a duration of 8000 ms. For Fig. 3 and later, the recurrent network was then trained with random initial conditions and noise amplitude 0.05 according to the algorithm described in equations (3-5). The fast target (2x speed) was trained over the window *t* ∈ [0,2000] with *y_SI_* = 0.3 and the slow target (0.5x speed) over the window *t* ∈ [0,8000] with *y_SI_* = 0.075. Ten differently seeded networks were each trained for a total of 60 trials alternating between fast and slow targets. A similar procedure was used to train networks at a single speed (Fig. 2). The initial target was captured with a duration of 4000 ms and *y_SI_* = 0.15 and zero noise. The same initial networks used in the temporal scaling task were trained at this speed for 30 trials. To suppress spontaneous activity, all networks were trained to maintain zero ***r*** (firing rate) for 30 s following the end of each trained recurrent target. We dubbed networks trained in this manner “gated” attractor networks because they only entered the long-lasting dynamic attractor in response to a specific cued input (Supplementary Fig. 1).

After recurrent training was complete, the output unit was trained, only at the fastest trained speed, to produce a target function of a series of 5 Gaussian peaks (“taps”) centered at 163, 513, 750, 1200, and 1750 ms (0.5x speed). The training algorithm for the output weights was similar to that used for recurrent training, as described previously^8^.

### Analysis of Temporal Scaling

To assess the ability of a network to generalize its activity to novel speeds, i.e. temporally scale, we tested the response of networks to a range of speed input levels after training was completed (weights were no longer modified). The network was set to a random initial state at *t* = −750 and given the trained cue input during *t* ∈ [−250,0]. The test speed input was delivered starting at *t* = −250 for a duration lasting 20% longer than when a perfectly timed last tap would occur. The timing of these peaks was used to measure the accuracy of the network’s temporal scaling using a *speed factor* and *scaling index*. The speed factor was a coarse measure of temporal scaling calculated by dividing the final peak time of twenty test trials at each speed to the mean peak time at the 1x speed, and taking the mean across trials. The scaling index was calculated by taking the mean fisher-transformed pairwise correlation of the mean timing of the five peaks for each speed against all other tested speeds

### Weber Analysis

The Weber analysis was performed according to Weber’s Generalized Law^51,52^, which defines a relationship between mean and variance of perceived time as:

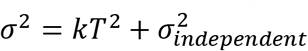

where *σ^2^* is the variance and *T* is mean time. We define the slope *k* as the Weber coefficient or slope;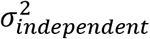 is the baseline (constant) variance sometimes referred to as motor variance. We measured *k* independently at each speed, by performing a linear fit on the measured mean and variance of the five response times at that speed (peak times for the RNN in Fig. 4 and button presses for psychophysics experiments in Fig. 5). Note that for visualization purposes, in some plots we show the linear fit of the standard deviation by *T*.

### Parallel Trajectories

To analyze the position of the trajectories in relationship to one another, we tested the networks at each speed without noise. We then concatenated the active period of the trajectory at each speed, defined as the window between cue input offset and speed input offset, and performed principal components analysis (PCA) on these concatenated trajectories. We used the PCA coefficients to transform the individual trajectory at each speed for visualization in Fig. 6a. To measure the relationship between trajectories, we returned to full (*N* = 1800) neural phase space and measured the Euclidean distance between the slowest (0.5x speed) trajectory and the trajectories at each speed, at all pairs of points in time. This produced one *t^test^* × *t^0 5x^* distance matrix per speed, as seen in Fig. 6b for test speed 2x. To confirm that the trajectories did not cross and followed a similar path, for each point on the slowest trajectory, we found a corresponding point on the test trajectory that was closest to it. This produced a vector of approximately 8000 distance values (for each millisecond of the slowest trajectory) which we plotted in Fig. 6c for each of the five tested speeds. The distances were relatively constant for each test speed and never reached zero, indicating that the trajectories did not intersect. We also recorded the points 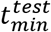 along the test trajectory where these minima occurred, allowing us to assess the relative speed of each trajectory along their entire length. For example, when the slowest trajectory is at its 400ms mark, if a test trajectory is closest to it at the test trajectory’s own 100ms mark, this would indicate that at that moment, the slowest trajectory was moving four times slower than the test trajectory. We plotted 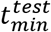 for each of the five tested speeds in Fig. 6d.

### Recurrent-Decay-Input Subspace Decomposition

In Fig. 7, the total drive (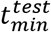, Equation l) was decomposed into its three components: 1) synaptic decay 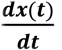; 2) recurrent synaptic drive 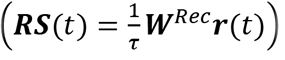; and its external component, the tonic speed input 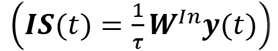. The magnitude of each of these components was calculated as the time-averaged L2-norm of the corresponding population vectors. Fig. 7C illustrates the generation of an orthonormal basis set {***is***, ***ds***, ***rs***} for the total drive at time *t*, which was computed by applying the Gram-Schmidt orthonormalization process as follows:

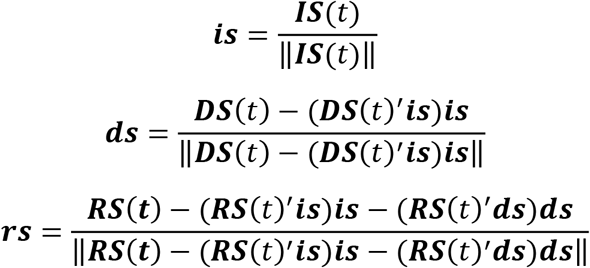

Here, ‖.‖ represents the L2-norm and the apostrophe represents the vector transpose operation. Collectively, these unit orthonormal vectors fully describe the total drive and its components at *t*, and therefore, form a basis set for these vectors. The plane described by the basis set {***ds***,***rs***} is denoted the internal drive plane, with ***DS***(*t*) projected onto this plane in grey, and ***RS***(*t*) in yellow. In Fig. 7D, we visualize the relationship between these vector projections over a short sequence of time steps along the slow and fast trajectories, on a common internal drive plane. For this, we constructed a common orthonormal set by applying the Gram-Schmidt process to the sequence-averaged component vectors. While doing so precludes the orthonormal set from forming a basis for the vector sequences, restricting the length of these sequences to a small fraction of the network unit time constant *τ*, renders the information loss negligible. Finally, in Fig. 7E, to show that the trajectories consistently encode their desired speeds, we plot the projection of the state variable (***x***(*t*)) onto ***is*** against its projection onto the first principal component in the subspace orthogonal to ***is***. That is, the x-axis represents the first principal component of (***x***(*t*) — (**x**(*t*)’ ***is***)***is***).

## Data and code availability

Data and code used to generate the results in this manuscript will be made available upon request, or code can be downloaded from:https://github.com/nhardy01/RNN

## Acknowledgements

This research was supported by NIH grants MH60163 and T32 NS058280, and NSF grant IIS-1420897.

## Author Contributions

D.V.B. conceived of the approach. N.F.H. performed the simulations and data analysis for the model. J.L.S., N.F.H. and D.V.B. designed the psychophysics experiments. J.L.S. conducted the psychophysics experiments and data collection. J.L.S. and D.V.B. performed data analysis for the psychophysics experiments. V.G. designed and performed the mechanisms experiments. N.F.H., V.G. and D.V.B. wrote the paper.

## Competing financial interests

The authors declare no competing financial interests.

